# miRNAs in platelet-poor blood plasma and purified RNA are highly stable: a confirmatory study

**DOI:** 10.1101/273797

**Authors:** Dillon C. Muth, Bonita H. Powell, Zezhou Zhao, Kenneth W. Witwer

**Author notes:** Corresponding author Kenneth W. Witwer, PhD 733 N. Broadway Miller Research Building Rm 829 Baltimore MD 21205 USA p 1-410-955-9770 f 1-410-955-9823. These authors contributed equally to this work (Dillon C. Muth, Bonita H. Powell).

## Abstract

The relative stability of microRNAs (miRNAs) as compared with other RNA molecules has been confirmed in many contexts. When bound to Argonaute (AGO) proteins, miRNAs are protected from degradation, even when released into the extracellular space in ribonucleoprotein complexes, and with or without the protection of membranes in extracellular vesicles (EVs). Purified miRNAs also appear to present less of a target for degradation than other RNAs. Although miRNAs are by no means immune to degradation, biological samples subjected to prolonged incubation at room temperature, multiple freeze/thaws, or collection in the presence of inhibitors like heparin, can typically be remediated or used directly for miRNA measurements. Here, we provide additional confirmation of early, well validated findings on miRNA stability and detectability. Our data also suggest that inadequate depletion of platelets from plasma may explain the occasional report that freeze-thaw cycles can adversely affect plasma miRNA levels. Overall, the repeated observation of miRNA stability is again confirmed.

## Introduction

microRNAs (miRNAs) are short (22 nucleotides on average) RNA molecules that contribute to post-transcriptional gene fine-tuning (1,2) and have aroused attention as potential biomarkers of disease (3–6) in part because of their stability. In cells, mature miRNAs are relatively long-lived, while messenger RNAs (for example) have short half-lives. A recent analysis of mRNA stability suggested that yeast mRNAs have a median half-life of only two minutes (7), even shorter than previously estimated. miRNAs seem to have greater stability, although longevity may differ by cell type and activity state (8,9). To provide just a few estimates of miRNA stability, published reports have placed miRNA half-life in the cell at: greater than eight hours (10); >24 hr, with “passenger” or “star” strands having substantially shorter half-lives (11); and around five days (12). In quiescent cells, miRNAs in inactive complexes can be stable for an astounding three weeks (13).

As recognized early on (14), mature miRNAs are more stable than other RNAs in biological matrices and after purification. miRNAs persist even in fixed, embedded tissue sections stored for years at room temperature. In biological fluids, similar persistence is observed, conferred by the tight AGO-miRNA association. In a landmark 2008 study, Mitchell, Parkin, and Kroh, et al observed that miRNAs were stable in plasma left at room temperature for up to 24 hours or frozen and thawed up to eight times (15). This observation has been confirmed repeatedly and extended to other biofluids and even longer storage times [see, for example (16–21)]. After purification, miRNA is also stable (22,23). When RNA samples were kept at near-boiling temperatures for up to four hours (24), MIQE-recommended RNA integrity numbers (RINs) (25) decreased over time; yet while mRNA detection declined, miRNA detection did not.

The stability of miRNAs in biological matrices is conferred by a tight association with Argonaute (AGO) proteins. Of two arms of a precursor miRNA, one is typically rapidly degraded, while the other is loaded during processing into an AGO (11). The AGO protects the mature miRNA for the remainder of its “life” and facilitates RNA regulation by bringing the miRNA and accessory proteins together in the RNA-induced silencing complex (RISC). Even outside the cell, miRNAs are found in AGO (26–28), whether within extracellular vesicles or not (29). Without this association, miRNAs are rapidly degraded in the RNase-rich biological compartments. Purified miRNAs (30) as well as synthetic miRNA sequences (15), unprotected by AGO, are degraded with a half-life of only seconds in room-temperature blood plasma.

Despite the near unanimity of observations on miRNA stability, there have also been several seemingly contradictory observations. Certain tissue-specific miRNAs in circulation, like miR-1 (muscle) or miR-122 (hepatocytes), have been reported to be more sensitive to freeze/thaw than others (31). Decreased detection of several plasma miRNAs after freeze/thaw cycles (32) or incubation of plasma at room temperature (33) was recently reported. To help resolve these apparent differences, we re-examined stability of several commonly investigated miRNAs in platelet-rich and platelet-poor plasma after incubation of plasma at 22 °C and after various freeze-thaw cycles. Since platelets—which are exquisitely temperature-sensitive and may be more susceptible to freeze-thaw damage than smaller carriers of miRNA—are not removed from plasma unless several centrifugations or other interventions are performed (34), we hypothesized that the profound influence of platelets may explain apparently contradictory findings in the literature.

## Methods

### Blood processing and plasma treatments

Fresh blood from human donors was obtained under a university-approved protocol (JHU IRB #CR00011400). Blood was collected into 60 mL syringes pre-loaded with 6 mL anticoagulant Acid Citrate Dextrose (ACD) (Sigma Aldrich, St. Louis, MO. Cat #: C3821. Lot #: SLBQ6570V). Whole blood was centrifuged within 15 minutes of draw at 1300 x g for 15 minutes to pellet blood cells. Supernatant (platelet-rich plasma or PRP) was aliquoted for later use or centrifuged twice at 2500 x g for 15 minutes. Supernatant from the final spin was defined as platelet-poor plasma (PPP). Three separate aliquots of both PRP and PPP were used for RNA extraction for each of the following conditions: fresh (immediate RNA isolation); one, two, three, four, five, and six freeze-thaw cycles (−80 °C to 22 °C), conducted regularly over four days and with rapid thaw; and 24-hour incubation at 22 °C.

### RNA extraction

RNA was extracted from 200 µl of plasma using the Exiqon Biofluids kit as described previously (35), including glycogen as carrier. RNA was stored at −80 °C until use unless otherwise specified.

### RNA stability experiment

Aliquots of purified plasma RNA were frozen at −80 °C, then thawed and incubated at room temperature for 7, 3, 1, and 0 days in a “countdown” design as previously described for shorter time periods (36). For the RNase A control, RNA was completely degraded by addition of equal volume RNase A (ThermoFisher, #EN0531, stock solution 10 mg/mL) prior to reverse transcription.

### qPCR assays

miRNA stem-loop reverse transcription quantitative PCR (37) was done as previously described (35) but using 384-well plates. Synthetic cel-miR-39-3p (Qiagen, #219610) was included in the reverse transcription master mix as a spike-in. Input into the reverse transcription reaction was normalized by volume (3 ul for plasma RNA). Assays were obtained from Applied Biosystems/Thermo Fisher, Cat. # 4427975: hsa-miR-16-5p (part number 000391), hsa-miR-21-5p (000397), and cel-miR-39-3p (000200). The quantitative step was done by following the manufacturer’s directions for cycling on a QuantStudio 12K Flex Real-Time PCR System (Thermo Fisher).

## Results

### Stability of miRNA in fresh vs frozen platelet-rich and –poor blood plasma

Whole blood samples were processed to platelet-rich plasma (PRP) immediately after draw. Platelet-poor plasma (PPP) was produced from a portion of the PPP. To assess effect of freeze-thaw cycle, PRP and PPP were frozen at −80 °C and thawed for a total of one through six cycles before RNA isolation. Triplicate aliquots were processed for each condition, and miRs-16-5p and −21-5p were measured by RT-qPCR. As shown in Figure 1, fresh PRP contained several fold more of both miRNAs than fresh PPP, with the difference attributable to the presence of platelets in the former. Compared with freshly processed plasma (“fresh”), a single freeze-thaw of PRP increased cycle numbers for detection of both miRNAs. Additional cycles of freeze-thaw had no consistent additional effect (Figure 1, FT 1-6). In contrast, miRNAs in PPP were insensitive to freeze-thaw. Although some technical variability was observed (see especially the outlier points FT3 for donor 1 plasma and FT1 for donor 2, Figure 1), the average Cq of miRNAs extracted from frozen/thawed PPP was indistinguishable from that of fresh plasma. There was no consistent trend towards higher Cq with higher numbers of freeze-thaw cycles.

**Figure 1.**
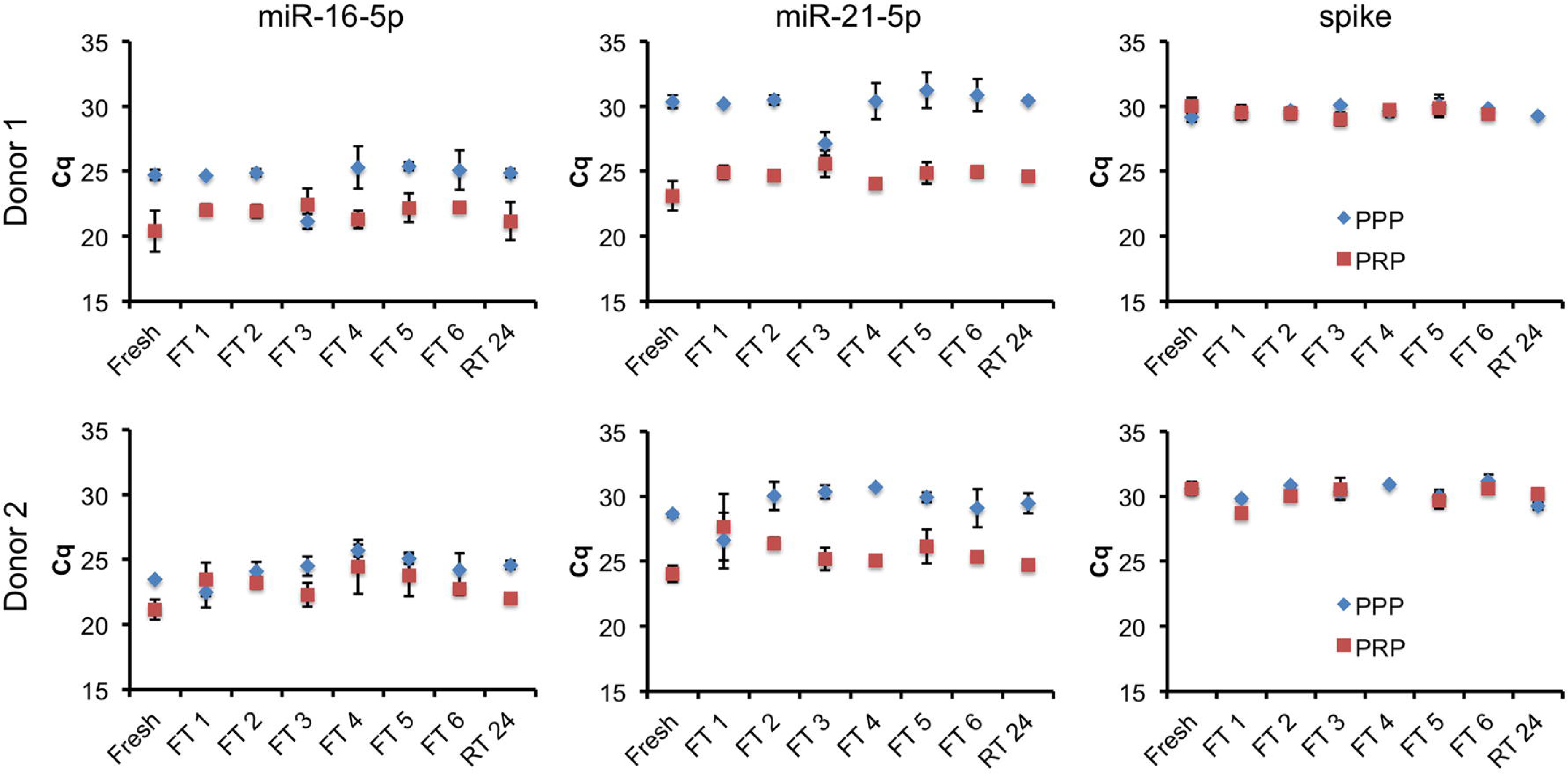
Stability of blood plasma miR-16-5p and miR-21-5p to freeze-thaw and room temperature incubation is associated with removal of platelets. Total RNA was isolated from platelet-rich plasma (PRP) or platelet-poor plasma (PPP) immediately after processing (Fresh), after one through six freeze-thaw cycles of −80 °C to 22 °C (FT 1 to 6), or after incubation of plasma at 22 °C for 24 hours (RT 24). miR-16-5p and miR-21-5p levels as well as a synthetic cel-miR-39-3p spike-in (spike) were assessed by stem-loop/hydrolysis probe qPCR assays, with results presented as Cq. Data are average plus and minus standard deviation for processing replicates. Three processing replicates and three qPCR measurements are included for each condition. No RT and no template reactions were also performed, with all Cq > 37 or undetected (not shown).

### Stability of miRNA in plasma to incubation at room temperature

In parallel with the freeze-thaw experiments, aliquots of PRP and PPP were incubated at room temperature (approximately 22 °C) for 24 hours before RNA isolation. PRP displayed a slight increase in Cq values for both miRNAs after this incubation, although not as pronounced as for the initial freeze-thaw cycle. Similar to the freeze-thaw results, miRNAs in PPP appeared to be insensitive to room temperature incubation (Figure 1, RT 24).

### Stability of miRNA in purified RNA

We have previously reported the stability of purified miRNAs in aqueous solution when incubated at room temperature for time periods ranging from 0 to 24 hours (36). We confirmed and extended this observation for miR-16-5p in total plasma RNA purified using the Exiqon Biofluids kit. RNA was incubated at 22 C for 0, 1, 3, and 7 days prior to qPCR. A slight increase in average Cq was observed after 7 days at room temperature (Figure 2). However, the range of values at day 7 overlapped with those from other days, and the difference was not significant. Addition of RNase A resulted in complete loss of signal.

**Figure 2.**
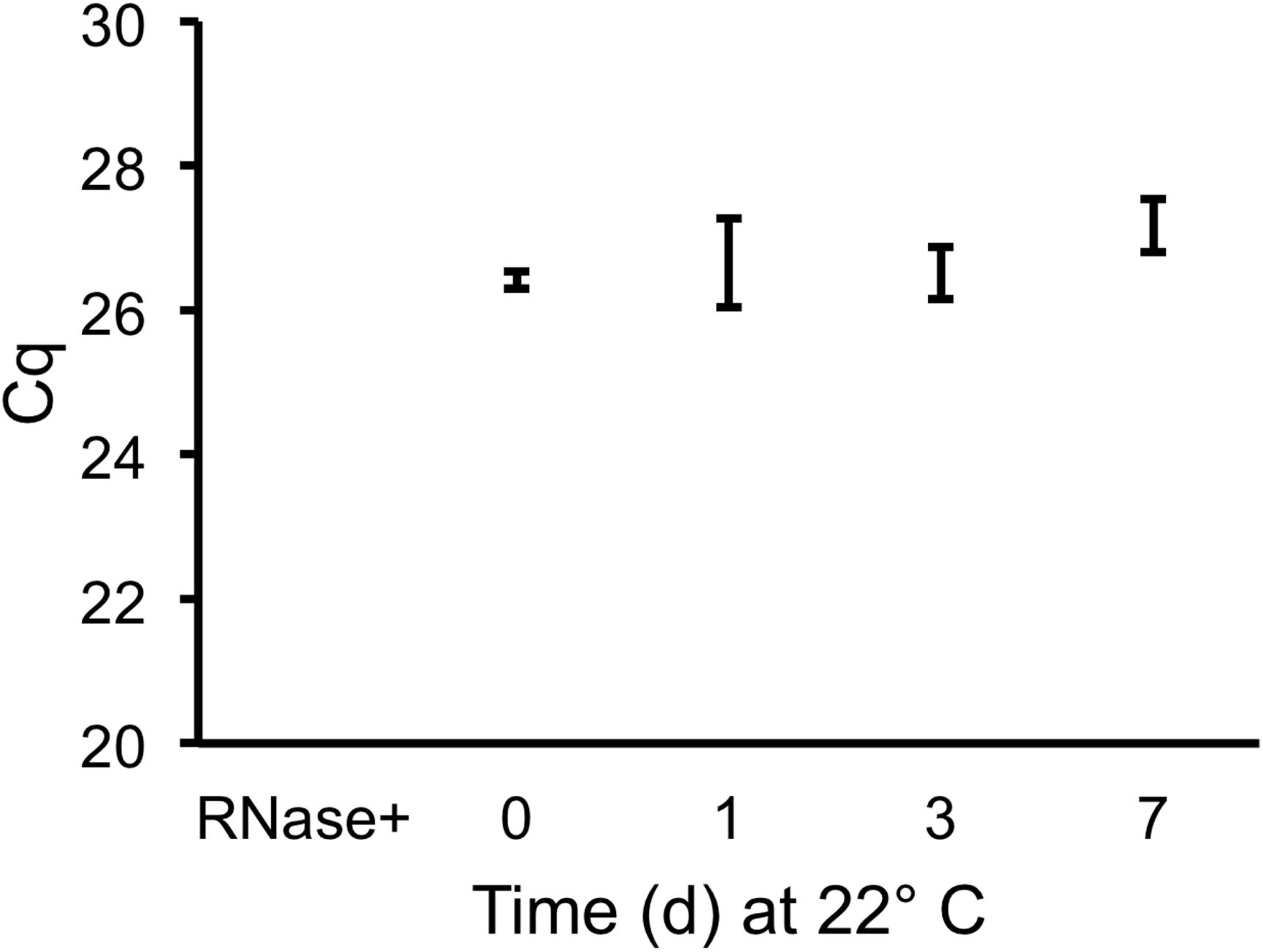
miR-16-5p in purified RNA is stable for up to one week at room temperature. Aliquots of total plasma RNA isolated from platelet-poor plasma (different batches from those shown in Figure 1) were frozen at −80 °C, then thawed and incubated at room temperature for 0, 1, 3, or 7 days, or thawed and treated with RNase A (Rnase+). qPCR for miR-16-5p was performed. Shown is average +/- standard deviation for three experiments.

## Conclusions

This study re-examined stability of miRNAs in blood plasma, probing abundance of miRNAs in PRP and PPP subjected to eight conditions including freeze-thaw (one to six cycles) and incubation at room temperature for 24 hours. Each condition was represented by three processing replicates (for a total of 96 plasma and RNA samples) and three qPCR measurements of RNA from each processing replicate. Like others, we observed no discernible, consistent effect of freeze-thaw or room temperature incubation on miRNA abundance in PPP. In contrast, miRNAs in PRP were affected by one freeze-thaw cycle and, possibly but to a lesser extent, by incubation at room temperature for 24 hours. However, freeze-thaw cycles after the initial cycle had no consistent effect on miRNA abundance even for PRP.

These results again reinforce that platelets contain the majority of miRNAs in platelet-rich plasma, and that platelets are more susceptible to a single freeze-thaw cycle or incubation at room temperature than smaller miRNA carriers such as exRNPs and EVs that remain after platelet removal. Once damage of platelets occurs in platelet-rich plasma, additional freeze-thaw cycles do not appear to exacerbate miRNA loss. While it is possible that different miRNAs would yield different results, this is most likely to occur in PRP, and for miRNAs with different abundance ratios in platelets versus true exRNA fractions. Studies that aim to examine truly extracellular plasma RNA must include careful separation of platelets and their RNA from other RNA sources.

Another potentially important conclusion is that small fold changes in extracellular miRNAs require multiple time points and replicates on which to base any firm conclusions. In this experiment, operators with multiple years of experience with RNA SOPs nevertheless observed anomalous results for several time points or conditions. These results could have led to incorrect conclusions if examined in isolation. For example, the “freeze-thaw 3” condition for PPP (donor 1) and the “freeze-thaw 1” condition for PPP (donor 2) appeared to show an increase in miRNA detection that was not seen for the spiked-in control. The differences we observed were clearly artifactual and introduced at some post-processing or RNA purification step, as a synthetic sequence spiked in just before qPCR was consistently and relatively invariantly detected for all conditions. Without the context of the other conditions/time points, these results might not have been identified as outliers and could have been incorrectly interpreted.

The remarkable stability of miRNAs, both in biological matrices and after RNA purification, has made miRNAs into attractive perceived biomarkers despite their comparative dearth of information content. Protected by a tight association with AGO proteins, mature miRNAs can be detected in biological matrices many years after tissue fixation or freezing, or after weeks or months at above-freezing temperatures, while miRNAs in purified RNA seem to present a smaller “target” for degradation and decay than other RNA species. To be sure, the best practice is always to compare directly only those samples that have been processed and treated in the same way. However, clinical samples and isolated RNA samples alike that have been stored for extended periods, left at room temperatures for hours to days, or subjected to multiple freeze-thaw cycles may still harbor miRNAs with an important tale to tell.

## ACKNOWLEDGEMENTS

The authors gratefully acknowledge Mr. J. Woodland Pomeroy for assistance with donor scheduling and the JHU DNA Analysis Facility for access to the QuantStudio qPCR system.

## AUTHOR CONTRIBUTIONS

All authors have accepted responsibility for the entire content of this submitted manuscript and approved submission. DCM, BHP, and ZZ collected samples and performed experiments and analyses; KWW planned and directed the studies, conducted analyses, and wrote the manuscript; all authors contributed to editing and revision of the manuscript.

